# Structure of an Hsp90-immunophilin complex reveals cochaperone recognition of the client-maturation state

**DOI:** 10.1101/2021.01.21.427690

**Authors:** Kanghyun Lee, Aye C. Thwin, Eric Tse, Stephanie N. Gates, Daniel R. Southworth

**Affiliations:** Department of Biochemistry and Biophysics, Institute for Neurodegenerative Diseases, University of California, San Francisco, CA 94158, USA; Graduate Program in Chemical Biology, University of Michigan, Ann Arbor, MI, USA

## Abstract

The Hsp90 chaperone promotes the folding and activation of hundreds of client proteins in the cell through an ATP-dependent conformational cycle guided by distinct cochaperone regulators. The FKBP51 immunophilin binds Hsp90 with its tetratricopeptide repeat (TPR) domain and catalyzes peptidyl-prolyl isomerase (PPIase) activity during the folding of kinases, nuclear receptors and tau. Here we have determined the cryo-EM structure of the human Hsp90:FKBP51:p23 complex to 3.3 Å that, together with mutagenesis and crosslinking analysis, reveals the basis for cochaperone binding to Hsp90 during client maturation. A helix extension in the TPR functions as a key recognition element, interacting across the Hsp90 C-terminal dimer interface presented in the closed, ATP conformation. The PPIase domain is positioned along the middle domain, adjacent Hsp90 client binding sites, while a single p23 makes stabilizing interactions with the N-terminal dimer. With this architecture, FKBP51 could thereby act on specific client residues presented during Hsp90-catalyzed remodeling.

## Introduction

Heat shock protein (Hsp) 90 is a universally conserved molecular chaperone that is broadly essential to the stability of the human proteome through the folding and activation of numerous client protein substrates such as nuclear hormone receptors (glucocorticoid and estrogen receptors), transcription factors (HSF1, p53, and OCT4) and kinases (BRAF, Cdk4, and ErbB2/Her2) (Schopf et al., 2017). Hsp90 is upregulated in cancer cells (Whitesell and Lindquist, 2005) and many oncoproteins, such as Bcr-Abl (An et al., 2000) and ErbB2/Her2 (Xu et al., 2001), require Hsp90 interaction. Thus, Hsp90-specific inhibitors, including geldanamycin and its derivatives, are of significant importance as anti-cancer therapeutics (Trepel et al., 2010). Additionally, Hsp90 plays key roles in neurodegenerative disease pathways including regulating Tau modification and folding (Shelton et al., 2017) and inhibiting α-synuclein aggregation (Daturpalli et al., 2013).

Hsp90 forms dynamic macromolecular assemblies with distinct cochaperone proteins that regulate its ATPase cycle and client folding steps (Pearl and Prodromou, 2006). Moreover, the interaction by the tetratricopeptide repeat (TPR) class of cochaperones is critical in conferring distinct Hsp90 folding steps and pathways, such as client loading (Hop), ubiquitination (Chip), and de-phosphorylation (PP5) (Biebl and Buchner, 2019; Smith, 2004). These cochaperones bind the EEVD C-terminal peptide of eukaryotic Hsp90s and Hsp70s via the 7-member α-helical TPR domain, which forms a conserved carboxylate “clamp” interaction with the terminal Asp (Scheufler et al., 2000). However, little is known about the structural basis for their interaction beyond TPR-EEVD binding or how distinct regulatory functions are specified on Hsp90.

The FK506-binding protein 51 (FKBP51), a member of the immunophilin family that includes FKBP52 and Cyp40 TPR-cochaperones, binds Hsp90 but not Hsp70 and catalyzes *cis-trans* peptidyl prolyl isomerization (PPI) of clients, regulating their folding and downstream function (Zgajnar et al., 2019). PPIase activity occurs through the N-terminal FK1 domain and is inhibited by immunosuppressive compounds including FK506 and rapamycin (Hahle et al., 2019). FK1 is connected to a related but inactive FK2 domain followed by the C-terminal TPR domain, which together adopt an extended tripartite configuration in the crystal structure (Sinars et al., 2003). Both FKBP51 and the close homolog, FKBP52, are identified in Hsp90:p23 maturation complexes containing the glucocorticoid or other nuclear receptor clients (Grad and Picard, 2007; Pratt and Toft, 1997), and have distinct functions in potentiating ligand binding and nuclear localization (Riggs et al., 2003; Storer et al., 2011; Wochnik et al., 2005). Notably, FKBP51 was recently discovered to act on Cdk4 in the Hsp90-stabilized complex, revealing a key role for its PPIase activity in Cdk4 inhibition (Ruiz-Estevez et al., 2018). Additionally, Hsp90:FKBP51 interacts with Tau, increasing stability (Jinwal et al., 2010) and promoting neurotoxic accumulation in neurodegenerative disease pathways (Blair et al., 2013).

How FKBP51 recognizes and acts on Hsp90-bound clients is unknown. Hsp90 exists as a homodimer through a C-terminal domain (CTD) interface and undergoes large, nucleotide-specific conformational changes during its chaperone cycle. The N-terminal ATPase domains (NTDs) dimerize upon ATP binding, forming a closed conformation that is activated for hydrolysis. The ability to form this closed, ATP state is rate limiting for hydrolysis (Prodromou et al., 2000; Richter et al., 2008) and varies across homologs (Southworth and Agard, 2008). Cochaperones Aha1 and p23 recognize the closed state and competitively bind NTD-MD sites across the dimer, thereby promoting (Aha1) or inhibiting (p23) ATPase activity (Ali et al., 2006; Meyer et al., 2004; Schopf et al., 2017), The structure of the closed state Hsp90:Cdc37:Cdk4 complex reveals the kinase substrate is dramatically unfolded with its N-lobe threaded between the monomers, contacting hydrophobic substrate-binding sites in the MD (Verba et al., 2016). While biochemical studies establish FKBP51 functions with p23 during client maturation (Ebong et al., 2016; Grad and Picard, 2007; Ni et al., 2010), recent NMR studies indicate FKBP51 stabilizes the Hsp90 open state to facilitate Tau binding (Oroz et al., 2018).

Here we sought to determine the structural basis for FKBP51 interaction with Hsp90 during the nucleotide-dependent chaperone cycle. Focusing on human FKBP51, p23 and Hsp90α, we developed *in vitro* conditions that promote and stabilize the closed, ATP conformation of Hsp90, enabling characterization of binding interactions between defined open and closed states. We find that FKBP51 preferentially binds the closed, ATP state, forming a complex with a 2:1 stoichiometry that is further stabilized by p23 binding. By cryo-EM we determined a 3.3 Å structure of the Hsp90:FKBP51:p23 complex that reveals distinct contacts by FKBP51 and define its asymmetric interaction and recognition of Hsp90. Photocrosslinking and mutagenesis further establish key interactions by the FK1 and TPR domains. From these results we propose a model in which helix 7 (H7) of the TPR domain functions as a critical specificity element for engaging Hsp90 in the p23-stabilized, ATP state to promote client maturation. This arrangement flexibly positions the FK1 domain adjacent Hsp90 client binding sites in a manner we predict facilitates site-specific PPIase activity towards clients in coordination with the Hsp90 chaperone cycle.

## Results

### FKBP51 preferentially recognizes the closed Hsp90 conformation

Immunophilins are present with p23 in Hsp90-steroid hormone receptor maturation complexes (Ebong et al., 2016; Johnson and Toft, 1994; Nair et al., 1997), indicating potential binding to the closed, ATP conformation of Hsp90 (Ali et al., 2006; Sullivan et al., 2002). While FKBP51 interaction with the Hsp90 open state has been characterized previously (Oroz et al., 2018), we hypothesized that this interaction may differ between the open and closed states. Therefore, we sought to characterize FKBP51 binding to the open (apo) and closed (ATP) states of Hsp90. Human Hsp90 highly favors the open conformation even when bound to the nonhydrolyzable ATP analog, AMPPNP, contrary to *E. coli* and yeast homologs (Southworth and Agard, 2008). Therefore, reaction conditions were investigated to stabilize the closed state of human Hsp90α. Based on the temperature dependence of the mitochondrial Hsp90 (TRAP1) closed state (Partridge et al., 2014), temperature (0°C, 25°C, and 37°C) and salt concentration (10, 50, 250, and 500 mM KCl) were tested in promoting the Hsp90α closed state with AMPPNP (Figure S1A). By native gel two distinct Hsp90 species are identified, with a faster-migrating band appearing following incubation with saturating (2 mM) AMPPNP (Prodromou et al., 1997) at 37°C for 2 hours compared to 0°C or without AMPPNP (Figure 1A). Notably, a complete shift to the faster-migrating band appears following the addition of high salt (500 mM), indicating both an increase in temperature and salt concentration with AMPPNP promotes formation of this species, which likely represents the closed conformation based on its higher mobility (Figure 1A). Only a minor amount of the higher mobility band is observed following incubation at 25°C, further supporting the high (37°C) temperature and salt dependence (Figure S1A). Finally, an increase in this species is observed with (NH_4_)_2_SO_4_ compared to NaCl and KCl, indicating salts that are more kosmotropic may be favorable (Figure S1B).

**Figure 1.**
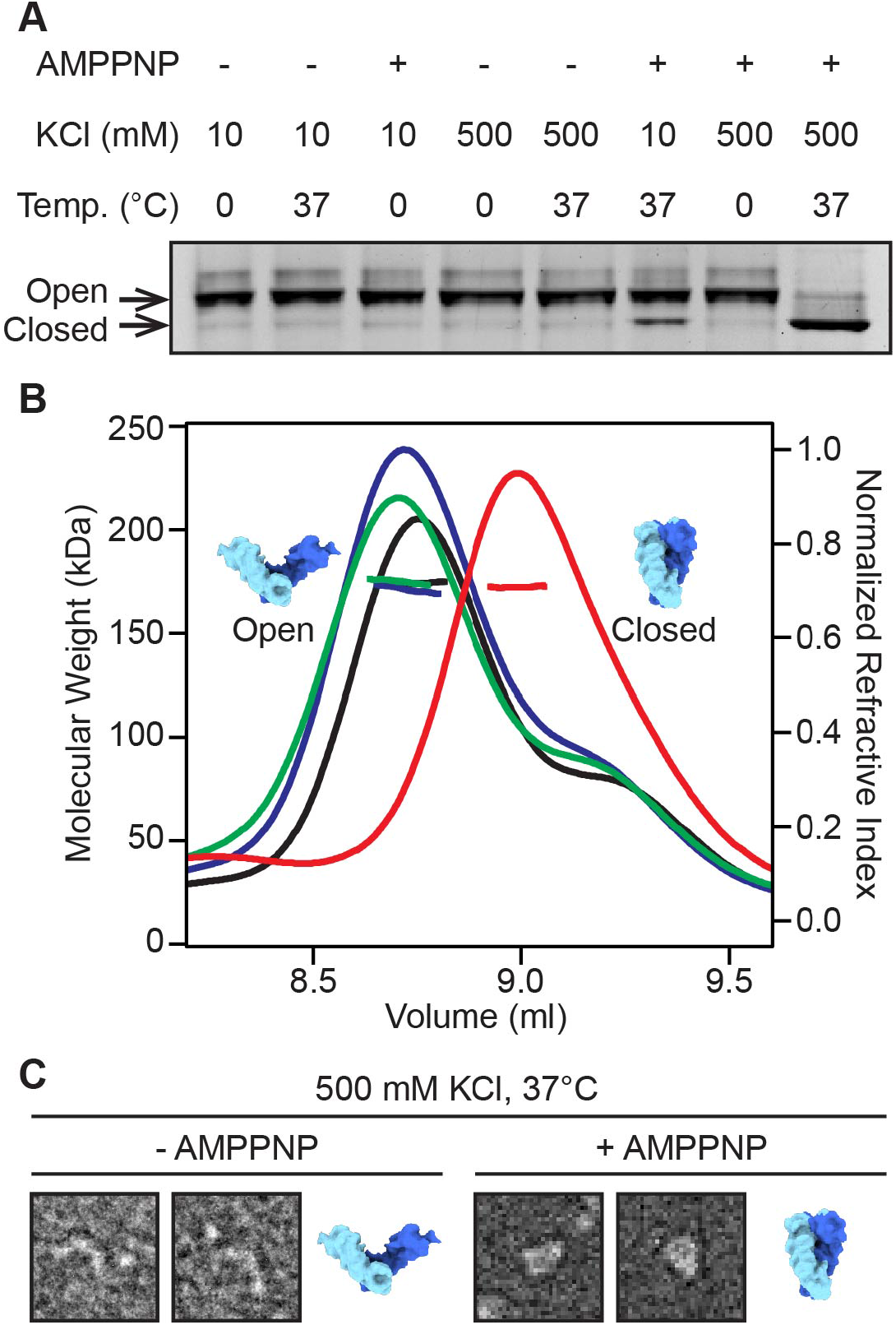
Analysis of the Hsp90 dimer in the open and closed states. (A) Native-gel separation of Hsp90 following incubation under indicated temperature, KCl and nucleotide (AMPPNP) conditions, showing shift to higher-mobility closed conformation at 37°C with 500 mM KCl and 2 mM AMPPNP. (B) SEC-MALS analysis of Hsp90 following 2 hours incubation at 37°C with no nucleotide and 500 mM KCl (black), at 0°C with 2 mM AMPPNP and 500 mM KCl (green), at 37°C with 2 mM AMPPNP and 8 mM KCl (blue), and at 37°C with 2 mM AMPPNP and 500 mM KCl (red). The mwavg determined by MALS is indicated by horizontal lines (kDa, left Y-axis), and is shown with the elution trace of the protein concentration (refractive index, right Y-axis) versus elution volume (ml). The shift to the later elution volume with no change in mwavg (red) indicates formation of the closed state. (C) Representative negative-stain single particle images of Hsp90 following incubation with 500 mM KCl at 37°C in the absence (left) and presence (right) of 2 mM AMPPNP and compared to structures of the Hsp90 dimer in the open (PDB: 2IOQ) and closed (PDB: 2CG9) states. See Figure S1C for more particle images.

Hsp90 conformations were additionally characterized by size-exclusion chromatography coupled to multiangle light scattering (SEC-MALS) (Figure 1B). Apo Hsp90 elutes at ∼8.8 ml (black) with an average molecular weight (mw_avg_) of 174 kDa, indicating a dimer species based on its calculated molecular weight (mw_calc_) of 169 kDa. Following incubation with AMPPNP at 37°C in high salt (500 mM KCl) the Hsp90 elution peak shifted to ∼9.0 ml (red), while the mw_avg_ remained similar (172 kDa), indicating a conformational change to a more compact dimer state. As with the native gel analysis, incubation at a lower temperature (green) or lower salt concentration (blue) resulted in only slight shifts in the elution profile, indicating both higher temperature and salt concentration promote this nucleotide-dependent conformational change. The Hsp90 conformation was verified directly by negative-stain EM (Figure 1C and S1C). In the absence of AMPPNP, Hsp90 particles adopt an open, extended conformation reflective of the apo state, similar to previous studies (Southworth and Agard, 2008). Conversely, analysis of Hsp90 incubated with AMPPNP and high salt at 37°C reveals the particles shift to a closed conformation which matches the NTD-dimerized, ATP state (Ali et al., 2006; Verba et al., 2016). Together, these results reveal that for nucleotide-bound human Hsp90α, high ionic strength and temperature together promote formation of the closed, ATP state.

Next, interaction between FKBP51 and Hsp90 in the open and closed states was characterized by SEC-MALS. Incubations were performed in the presence of 500 mM KCl at 37°C with and without AMPPNP. For Hsp90 incubated with FKBP51 in the absence of nucleotide the predominant peak elutes at ∼8.8 ml with a mw_avg_ of 180 kDa which is slightly higher than apo Hsp90 alone, indicating potentially weak binding of FKBP51 (Figure 2A). Strikingly, incubation of Hsp90 and FKBP51 with AMPPNP under the closed state conditions resulted in peak elution at ∼8.9 ml and a mw_avg_ of 218 kDa (Figure 2A). This is a shift by ∼0.1 ml and an increase in mw_avg_ by 46 kDa compared to the closed state Hsp90, indicating formation of an Hsp90:FKBP51 complex with an approximately 2:1 stoichiometry, based on the mw_calc_ of 51 kDa for FKBP51. Analysis of the peak fractions revealed Hsp90 and FKBP51 co-elute in this peak, with AMPPNP, while a minor amount of FKBP51 is present in the absence of AMPPNP (Figure S2A and S2B). These results therefore indicate that while lower-affinity interactions may occur with the open state of Hsp90, FKBP51 more stably binds the closed, ATP conformation of Hsp90, thereby forming a 2:1 asymmetric complex. not change substantially following incubation with p23, while a modest increase in the mw_avg_ to 183 kDa was identified under closed-state conditions with AMPPNP, indicating partial p23 binding (Figure 2B). By gel analysis of fractions from SEC, p23 co-elutes with the closed state Hsp90 but is not present in the apo Hsp90 fractions, indicating preferential binding to the closed state (Figure S2A and S2C). Thus, while the interaction appears to be low affinity with a possible 2:1 stoichiometry, this supports the known p23-specificity for the closed, ATP state of Hsp90 (Richter et al., 2004; Siligardi et al., 2004). When p23 was next incubated with Hsp90 and FKBP51 under the closed-state conditions the mw_avg_ increased to 225 compared to 172 and 218 kDa for Hsp90 alone or with FKBP51, respectively (Figure 2C). Both p23 and FKBP51 co-elute with the closed-state Hsp90 in these fractions, indicating formation of a ternary complex with FKBP51 (Figure S2A and S2D). No substantial changes in elution or mw_avg_ were observed under apo-state conditions compared to Hsp90 incubated with FKBP51 (Figure 2C). Together these results indicate that FKBP51 preferentially recognizes the closed, ATP state of Hsp90, binding asymmetrically with a 2:1 stoichiometry. Moreover, this interaction is compatible with p23 binding, resulting in the formation of a ternary, client-maturation complex.

**Figure 2.**
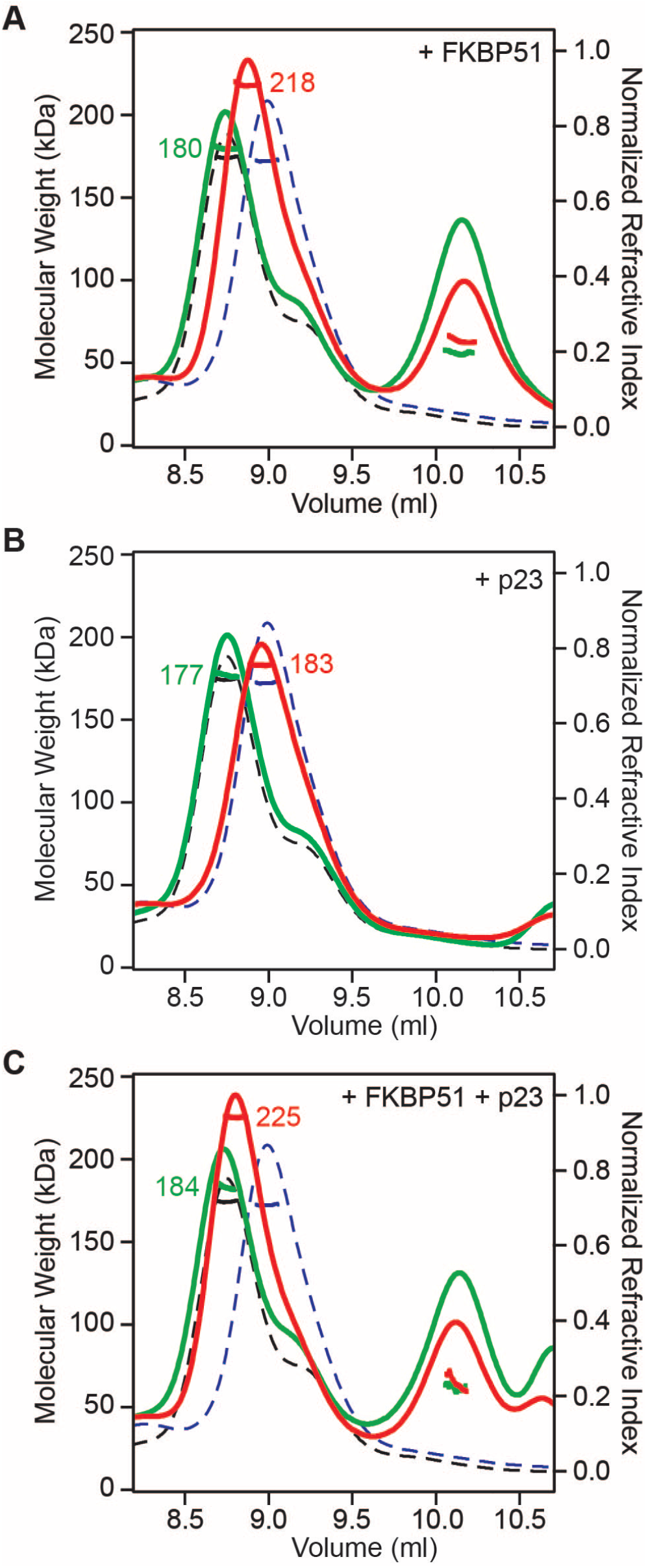
SEC-MALS analysis of FKBP51 and p23 binding to Hsp90 in the open and closed states. (A) FKBP51, (B) p23, or (C) FKBP51 with p23 were incubated with Hsp90 at 37°C in 500 mM KCl under open-state (green) or closed-stte (with 2 mM AMPPNP) (red) conditions for 2 hours and compared to Hsp90 alone incubated under the same conditions for the open (black, dashed) and closed (blue, dashed) states. Axes are shown as in Figure 1 and the mwavg (kDa), determined by MALS, is shown for the corresponding Hsp90 complex.

### Cryo-EM structure of the Hsp90:FKBP51:p23 complex

We next performed cryo-EM to determine the structural basis of the Hsp90:FKBP51:p23 interaction. Initial screening by cryo-EM indicated the complex is unstable, likely due to the vitrification conditions. Therefore, in order to improve stability and occupancy of FKBP51, samples were crosslinked by low-level (0.01%) glutaraldehyde crosslinking (Southworth and Agard, 2011). The 2D class averages of crosslinked Hsp90:FKBP51:p23 appear homogenous, exhibiting the closed-state architecture of Hsp90, but with additional density extending from CTD region (Figure 3A and S3A).

**Figure 3.**
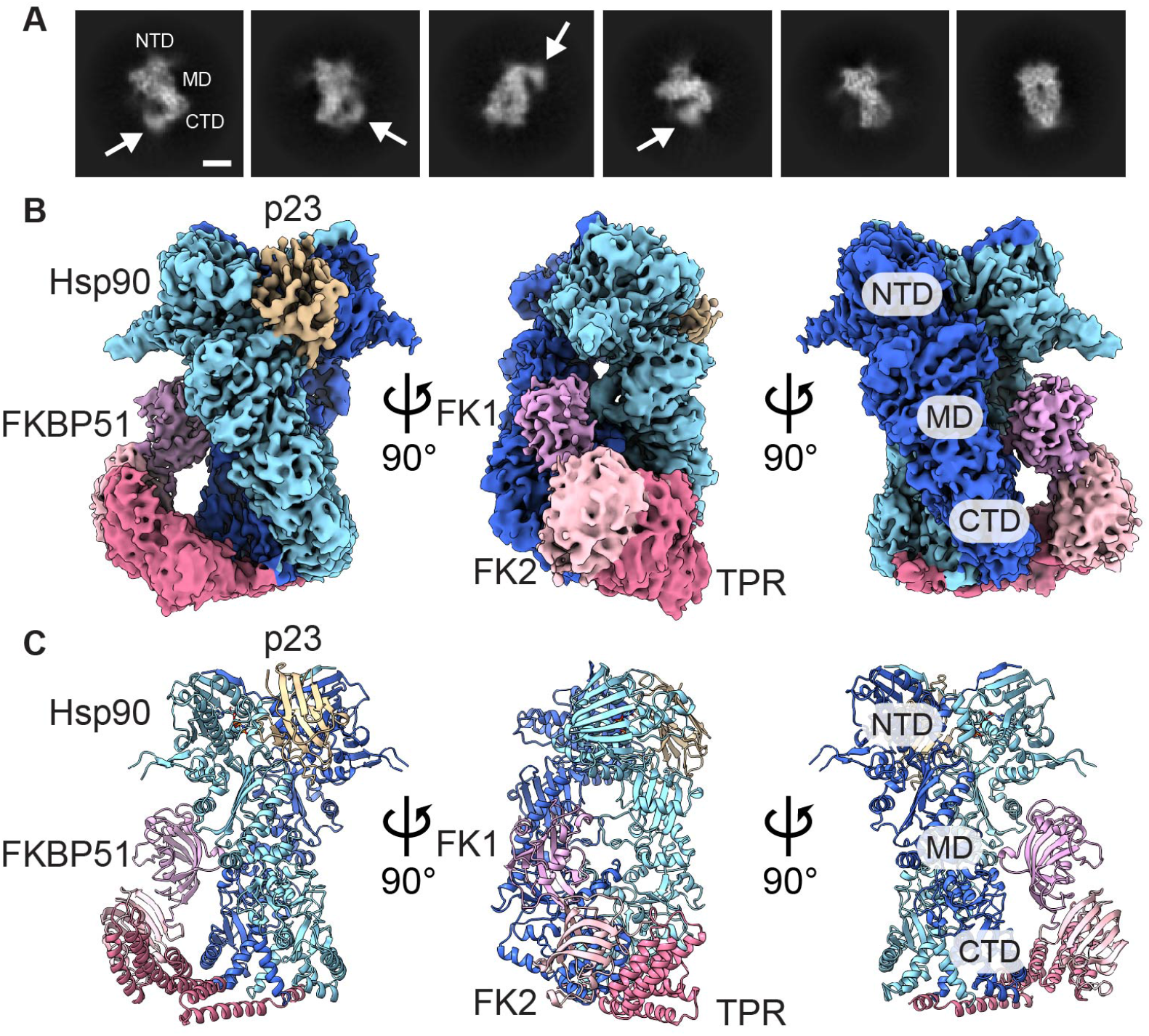
Cryo-EM structure of Hsp90:FKBP51:p23 closed-state complex. (A) Representative 2D class averages of Hsp90:FKBP51:p23 (scale bar = 50 Å). The Hsp90 domains are indicated and additional density corresponding to FKBP51 is shown (arrow) adjacent the CTD, based on comparison to the Hsp90 closed state structure. See Figure S3A for more 2D class averages. (B) The final cryo-EM map of Hsp90:FKBP51:p23. Densities are colored based on the molecular model and correspond to the Hsp90 monomers (light and dark blue); the FKBP51 domains: FK1 (plum), FK2 (light pink) and the TPR (dark pink); and p23 (tan). (C) The final molecular model of Hsp90:FKBP51:p23 colored as in (B).

Following 3D classification and refinement a final map at 3.3 Å overall resolution was achieved containing well-resolved density for the Hsp90 dimer and singly-bound FKBP51 and p23, supporting the asymmetric 2:1:1 arrangement (Figure 3B, S3B, S3C and Movie S1). Particle orientations are well-distributed with some preferred side-views (Figure S3D). Hsp90 adopts the canonical two-fold symmetric closed state conformation in which dimeric interactions occur in both NTDs and CTDs. The Hsp90 dimer is at a higher resolution (< 3.0 Å) while density for FKBP51 and p23 is lower (5-8 Å and 3-6 Å, respectively), indicating flexibility or reduced occupancy (Figure S3E). Nonetheless, density for all three FKBP51 domains is identified, with strong density for the TPR α-helical bundle adjacent the Hsp90 CTD and weaker density for the FK1 and FK2 domains present along the MD of one monomer (Figure 3B). Density corresponding to one p23 molecule is present on the opposite side of Hsp90, interacting at the cleft between the two NTDs (Figure 3B).

The reconstruction was sufficient to build a full atomic model of the Hsp90:FKBP51:p23 ternary complex by rigid-body docking homology models from existing structures and refinement with Rosetta (Figure 3C and S3F) (Kumar et al., 2017; Song et al., 2013; Verba et al., 2016; Weaver et al., 2000). The model for the Hsp90 dimer is similar to previous structures of yeast (Ali et al., 2006) and human (Verba et al., 2016) Hsp90 in the closed, ATP state (Cα-r.m.s.d. = 1.6 Å and 1.1 Å, respectively). NTD dimerization is defined by symmetric β-strand strap interactions across the top of the NTDs (residues 17-25), and the subsequent αH1 segment, which interacts across the dimer interface (Figure S4A). Similar to other ATP-state structures, the α-helical lid segment (residues 109-139) closes across the nucleotide pocket in both monomers and R400 in the MD contacts γ-phosphate of AMPPNP, forming the canonical Arg-finger interaction, which stabilizes the closed conformation required for hydrolysis (Figure S4B) (Ali et al., 2006; Cunningham et al., 2012). Additionally, MD amphipathic loops (residues 349-359 and residues 617-621) involved in substrate binding protrude into the dimer cleft (Figure S4C), and the CTDs interact primarily via hydrophobic contacts by the last two α-helices (αH17 and αH18, residues 657-695), which are essential for dimerization and hydrolysis (Figure S4D) (Minami et al., 1994; Prodromou et al., 2000).

FKBP51 fits unambiguously into the density and adopts a similar overall conformation as the crystal structure except for rotations around the FK2-TPR linker (Figure 3B and 3C) (Kumar et al., 2017). The majority of interaction with Hsp90 occurs through the TPR domain, which makes multiple surface contacts with the CTD (Figure 3C, discussed below). Notably, the FK1 domain makes minor contact with Hsp90 along the MD adjacent the substrate-binding loops. While p23 interacts with Hsp90 NTDs on the opposite dimer interface, the position of FKBP51 does not appear to overlap directly with where p23 could bind on the same side. Indeed, we identify partial density for p23 on the same side of Hsp90 as FKBP51 in one of the 3D classes (Figure S4E). Thus, p23 may be able to interact adjacent FKBP51 on the same side in certain client maturation complexes.

Another 3D class exhibited well-defined density for a single p23 but no FKBP51 (Figure S3B). This class refined to 3.1 Å and yielded improved density for p23 compared to the full complex (Figure S3C-F). The atomic model has a similar architecture as yeast Hsp90:p23 (Ali et al., 2006) and the recent structure of human GR:Hsp90:p23 (Noddings et al., 2020) (Figure S4F-I). Key interactions include contacts by conserved hydrophobic residues F103 and W106 of p23 with a hydrophobic pocket (residues L335, L394, P395, I408 and V411) in the MD of Hsp90 (Figure S4G) (Ali et al., 2006). K95 of p23 appears to form a salt bridge with E336 in the MD of Hsp90 (Figure S4F). However, other electrostatic interactions between p23 and Hsp90, including D122-K27 and K123-D193, identified in the yeast structure, are not conserved (Figure S4I). Alternatively, conserved residues R71 and D70 of p23 interact with Hsp90 S31, A166 and S165 (Figure S4H) and N97 (P115 in yeast) contacts M199, L122 and N123 in the lid of Hsp90, likely stabilizing the closed conformation (Figure S4F).

### Hsp90:FKBP51 interactions are defined by contact between the Hsp90 CTD and the TPR helix 7 extension

In the Hsp90:FKBP51:p23 structure the TPR domain of FKBP51 is oriented with the MEEVD binding pocket positioned away from the Hsp90 CTD (Figure 4A). Additional density is identified in this pocket and a DTSRMEEVD peptide docks appropriately into the density based on previous crystal structures (Figure 4A) (Kumar et al., 2017). However, the resolution was not sufficient to confirm the sequence. This peptide connects to the Hsp90 CTD by a ∼30-residue linker which is not visible, but likely sufficient to span the ∼40 Å needed to reach the last C-terminal residues (I698) resolved in the structure at the base of the CTD (Figure 4A, right). Thus, FKBP51 appears engaged with one MEEVD of Hsp90, however, the peptide may be derived from either monomer.

**Figure 4.**
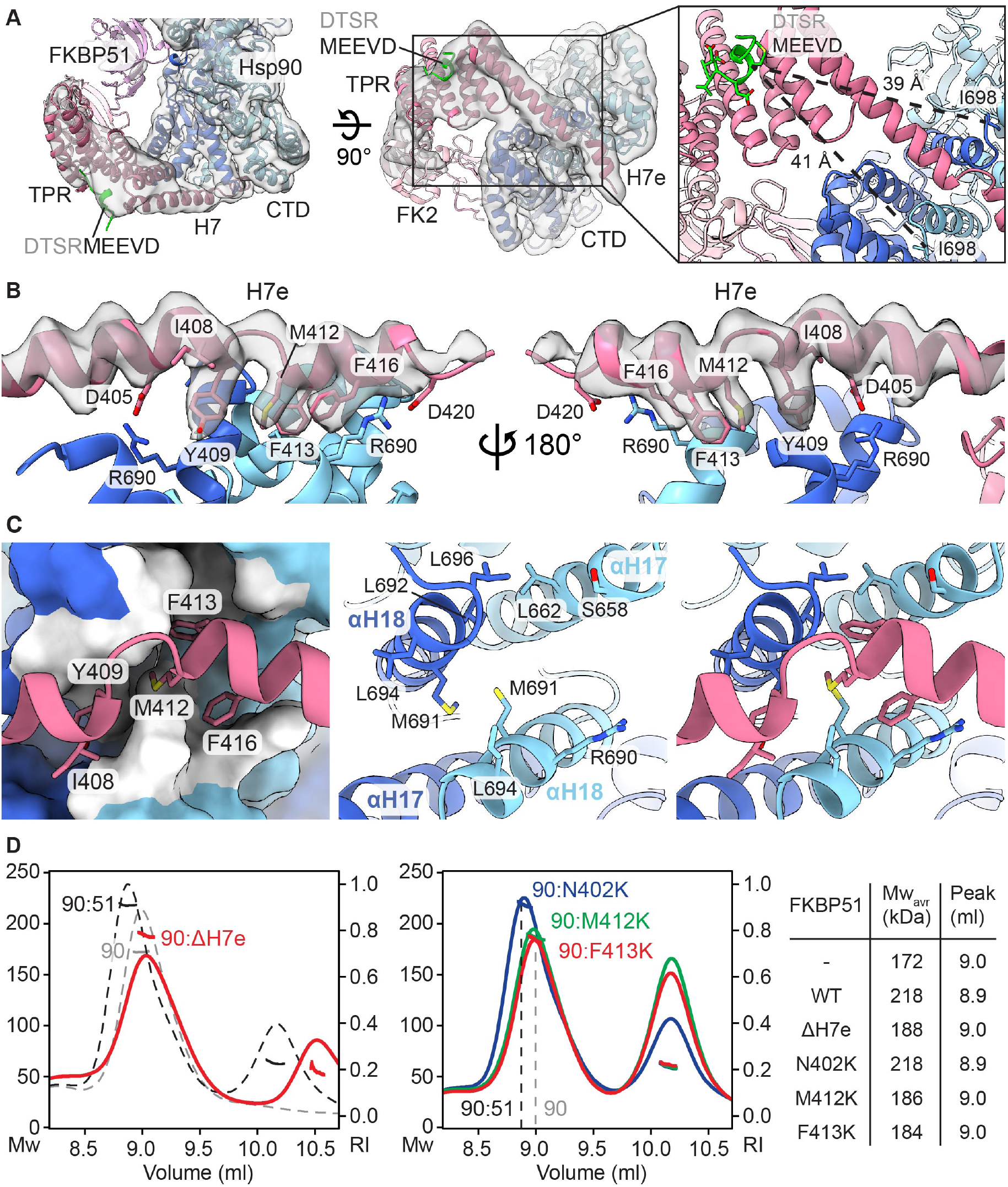
Contacts between FKBP51 TPR domain helix 7 extension (H7e) and the Hsp90 CTD define the closed state interaction. (A) Low-pass filtered cryo-EM density map and model of Hsp90:FKBP51:p23 showing the CTD-TPR interaction, colored as in Figure 3 with the Hsp90 MEEVD peptide (green) modeled based on the FKBP51 crystal structure (PDB: 5NJX) (Kumar et al., 2017). Expanded view (right panel) shows the approximate distances between M728 from the MEEVD and the Hsp90 C-terminal residue, I698, identified in the structure (dashed lines). (B) View of H7e (map+model) interacting across the Hsp90 CTD dimer interface. Proposed H7e hydrophobic interacting residues are labeled along with D405 and D420 residues which contact Hsp90 R690. (C) Surface representation of the Hsp90 CTD dimer groove is shown with hydrophobic residues in white and ribbon view of H7e (left panel); view of Hsp90 CTD dimer interface helices αH17 and αH18 and proposed interacting residues shown without (middle) and with H7e (right). (D) SEC-MALS analysis as in Figure 2 of Hsp90:FKBP51 complex formation under closed state conditions with FKBP51 variants: (left) ΔH7e (Δ401-457) (red), and (middle) M412K (green), F413K (red), and N402K (blue) in comparison to wildtype closed state complexes of Hsp90 (grey) and Hsp90:FKBP51 (black). The mwavg and peak elution volume are shown for each condition (right).

Surprisingly, the C-terminal H7 extension (H7e) of the FKBP51 TPR forms a distinct kinked conformation compared to crystal structures (Kumar et al., 2017; Sinars et al., 2003), and interacts directly with the Hsp90 CTD dimer (Figure 4B and S5A-C). H7e projects across the base of Hsp90, contacting both monomers in a hydrophobic cleft at the dimer interface comprised of the last two CTD helices (αH17 and αH18) (Figure 4B). Notably, the H7e helix residues 409-413 are distorted, enabling H7e to match the bend in the CTD cleft at the two-fold axis and contact both monomers (Figure S5A-C). Overall, the H7e region buries ∼830 Å^2^ of CTD surface area, which is larger than what is estimated for MEEVD interaction (∼500 Å^2^) (Kumar et al., 2017), thus revealing a new and substantial binding interface beyond the canonical TPR interaction.

This helix is well-resolved and distinct side-chain contacts with Hsp90 are identified (Figure 4B). Hydrophobic residues I408, Y409, M412, F413 and F416 in the unfolded strand of H7e interact with Hsp90 CTD residues 691-696, which together form distinct hydrophobic cleft between the C-terminal helices (αH17 and αH18) of each Hsp90 monomer (Figure 4C). Flanking this interaction, R690 from each Hsp90 monomer appear to make salt-bridge contacts with D405 and D420 in H7e on either side of the hydrophobic residues (Figure 4B). Additional contacts with Hsp90 occur between the H5-H6 connecting strand in FKBP51, including N365, which appears to make backbone contact with N655 in Hsp90 (Figure S5D). Overall, the H7e-CTD interaction appears to be a significant component of the Hsp90:FKBP51 maturation complex and the position enables H7e to make symmetrical contact across the CTD dimer, thereby precluding the binding of two FKBP51 molecules.

Next, we sought to characterize the significance of H7e in Hsp90 binding by mutagenesis and SEC-MALS analysis. Truncation of H7e from position H401 (ΔH7e) substantially reduced FKBP51 binding to Hsp90 under closed-state conditions, as indicated by the decrease mw_avg_ from 218 kDa to 188 kDa and shift in elution volume (Figure 4D). No substantial changes were observed under Hsp90 open-state conditions; both wt and ΔH7e exhibit incomplete binding to apo Hsp90 (Figure S5E). To test the significance of the hydrophobic interactions with the CTD cleft, mutations M412K and F413K were tested. N402K was included as a control due to its position outside the CTD interaction site, based on the structure. For M412K and F413K, the mw_avg_ is substantially reduced (186 kDa and 184 kDa, respectively) compared to the wt complex, and the elution volume overlaps with Hsp90 alone, indicating loss of FKBP51 binding (Figure 4D). The N402K mutation exhibits no significant change in the mw_avg_ and elution volume compared to the wt complex, validating that this site does not participate in Hsp90 binding. Together, these results establish that H7e is critical for FKBP51 binding to Hsp90. Moreover, the hydrophobic contacts with the Hsp90 C-terminal helices identified in the structure, involving the 408-416 unfolded strand, indeed appear to coordinate this interaction. The H7e itself, as well as residues Y409, M412, and F413 are conserved among the immunophilin-class of TPR cochaperones (Figure S5F). Thus, the interaction identified here likely represents a conserved Hsp90-specific recognition mechanism for these cochaperones.

### H7e binding to the CTD dimer groove likely specifies recognition of the Hsp90 closed state

Based on the SEC-MALS data (Figure 2) we identified that FKBP51 preferentially binds Hsp90 in the closed, ATP conformation compared open, apo state. However, FKBP51 does not appear to make substantial contact with NTD or MD regions that exhibit significant structural changes between these states (Figure 5A). Therefore, in order to understand this specificity, the CTD dimer conformation was compared between the open and closed states, based on structures of apo Hsp90 (*E. coli* HtpG) and Hsp90:Cdc37:Cdk4 in the closed state (Shiau et al., 2006; Verba et al., 2016). Notably, Hsp90 structure and sequence is highly conserved across species (39% sequence identity and 59% sequence similarity between *E. coli* HtpG and human Hsp90α). By global alignment (Cα) the CTD conformation does not vary substantially between the open and closed states (r.m.s.d = 1.8 Å) (Figure S6A). However, comparison of the CTD dimer interface reveals CT-helices αH17 and αH18 undergo distinct conformational changes that widen the CTD cleft from ∼6 to 12Å, thereby enabling the hydrophobic groove to become more accessible in the closed state (Figure 5A and Movie S2). By modeling H7e across the CTD in the apo state, residues M412 and F413 appear to clash with Hsp90 residues L662, M691, and L694, indicating the CTD may be incompatible for binding in this state (Figure 4C and 5B). In addition, changes in the CTD would shift αH17, bringing it closer to TPR helices H5 and H6 of FKBP51, likely altering interactions with FKBP51 N365 (Figure 5B and S5D). Based on these comparisons, the Hsp90 CTD dimer interface involving αH17 and αH18 undergoes conformational changes from the open to closed states that likely result in improved accommodation of the H7e strand, thereby specifying FKBP51-binding to the closed state of Hsp90.

**Figure 5.**
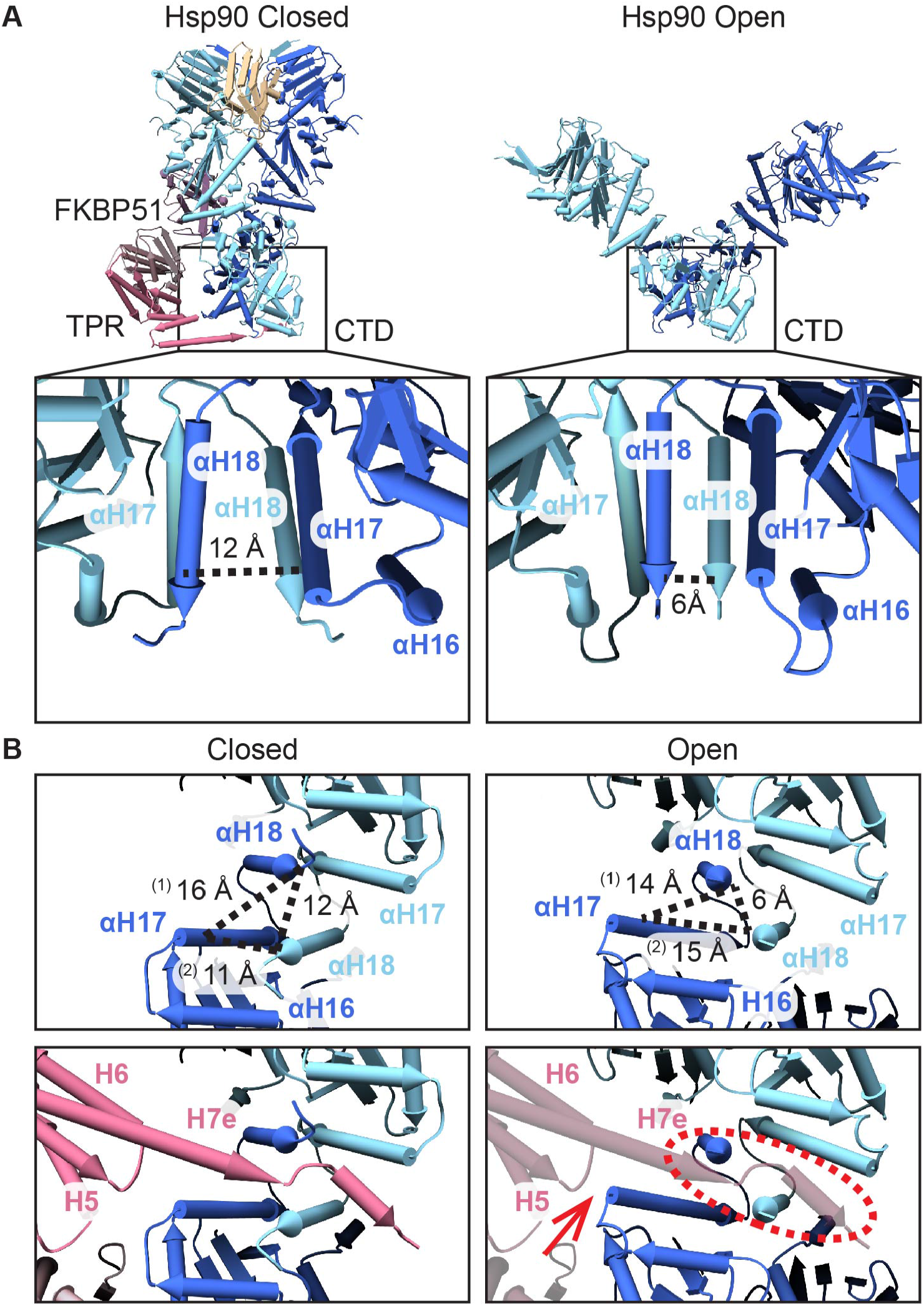
Comparison of Hsp90 CTD conformations showing accommodation of FKBP51 H7e is specific to the closed state. (A) Closed-(Hsp90:FKBP51:p23) and open-state (E. coli HtpG, PDB: 2IOQ) (Ali et al., 2006) Hsp90 structures with an expanded view of the CTD dimer showing the cleft between the monomers, at αH17 and αH18, widens towards the C terminus in the closed state. Distances between αH18, at I692, across the monomers is shown for the two states (dashed line). (B) Rotated end on view of the CTD dimer in the closed (left) and open (right) states showing distances between αH17 and αH18 across the monomers at indicated positions (dashed lines). Measurements are between αH17 (S658) and αH18 (I692) in the same subunit (1), and between αH17 (S658) and αH18 (I692) across subunits (2). FKBP51 TPR (pink) is shown with H7e bound across the Hsp90 CTD dimer in the closed state (lower, left panel), as modeled in the Hsp90:FKBP51:p23 structure, but is incompatible and clashes with the CTD in the open state (lower right panel, dashed circle). Additional clashing at the H5 loop is indicated (arrow).

### Position of the FK1 domain supports PPIase-directed activity on Hsp90-bound clients

In the Hsp90:FKBP51:p23 structure FKBP51 extends from the CTD-TPR interaction along the Hsp90 MD, enabling the N-terminal FK1 domain to be positioned adjacent Hsp90 client interaction sites (Figure 6A, bottom) (Genest et al., 2013; Verba et al., 2016). The FK1 PPIase active site is comprised of a series of β strands that form a conserved hydrophobic pocket where immunosuppressive drugs bind, including FK506 and rapamycin (Hahle et al., 2019). This pocket is approximately 17 Å from Hsp90 client binding sites, which include Y528, Q531, and M610 (Genest et al., 2013), and have been identified to contact the unfolded Cdk4 client in the Hsp90:Cdc37:Cdk4 closed state structure (Verba et al., 2016). In order to further characterize the proximity of the FK1 domain to Hsp90, site specific crosslinking was performed by incorporating the photoactivatable unnatural amino acid, p-benzoyl-l-phenylalanine (Bpa) into specific positions in FKBP51. A bulge within the β3 strand is the closest to the Hsp90 client biding site (∼5 Å away) and was thus chosen for Bpa incorporation (Figure 6A, top). Bpa was incorporated into D72, N74, and E75 and full-length protein was purified and crosslinked with Hsp90 under closed and open-state conditions (Figure 6B). By SDS-PAGE, distinct high-mw bands are identified for all 3 sites only in the presence of Hsp90 and following UV exposure, confirming that FKBP51 β3 is indeed in close proximity to Hsp90. Notably for D72Bpa, strong crosslinked bands are only present under the Hsp90-closed state condition, indicating that crosslinking at this position is conformation-specific. Additionally, N74Bpa and E75Bpa each show crosslinked-bands under open-state conditions, however, these bands diminish for the closed state, while several other bands appear, revealing distinct interactions in the open and closed states. Thus, Bpa incorporation at this β3-strand bulge position provides a crosslinking-readout for the Hsp90 conformation and confirms the FKBP51-specificity for the Hsp90 closed state and positioning of FK1 PPIase site adjacent Hsp90. Based on the close proximity of the PPIase site to the Hsp90 client binding strands we predict that FKPB51 may target specific client proline residues near where Hsp90 binds clients. Furthermore, the position of the β3-strand bulge is intriguing and may provide an important interaction link with Hsp90 to facilitate client positioning for PPIase activity.

**Figure 6.**
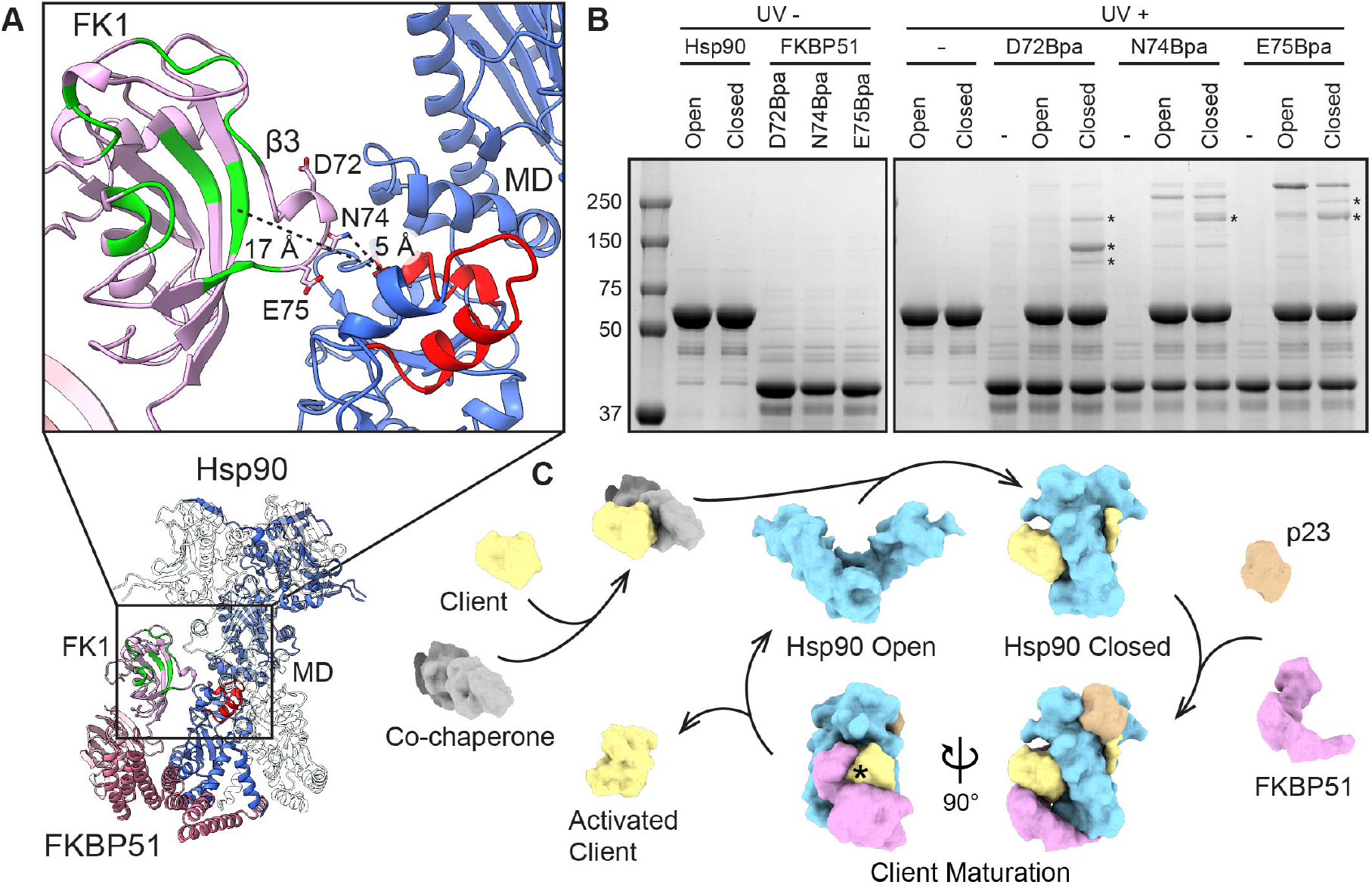
Positioning of FK1 PPIase domain adjacent Hsp90 client binding sites and model for FKBP51 function during client maturation. (A) The Hsp90:FKBP51:p23 structure with distances showing the FK1 PPIase site and FK506 binding pocket (green) and connecting β3 bulge are positioned adjacent MD client binding sites (red) in Hsp90. Sites of Bpa incorporation in the β3 bulge region are shown. (B) SDS-PAGE analysis of Hsp90 and FKBP51 variants containing Bpa at D72, N74, or E75 following incubation under closed- and open-state conditions and UV photocrosslinking. Unique crosslinking bands present under closed-state conditions are indicated (asterisks). (C) Model for Hsp90-catalyzed client maturation and activation based on the Hsp90:FKBP51:p23 structure in which FKBP51 binds Hsp90 with p23 following ATP binding and NTD dimerization to form the client maturation complex. In this arrangement, FKBP51 is positioned for PPIase activity directed at specific client sites presented through Hsp90 binding and remodeling, thereby promoting downstream client folding and activation.

## Discussion

The FKBP51 immunophilin serves critical roles in the Hsp90 chaperone pathway through PPIase pro-folding activity directed towards diverse Hsp90-bound clients including nuclear hormone receptors, kinases, and tau. FKBP51 has been proposed to act during client maturation, but the structural basis for how the TPR-containing cochaperone recognizes Hsp90 or is positioned for client interactions has been unclear. Here we identified that FKBP51 preferentially binds Hsp90 in the closed, ATP state and determined a structure of the Hsp90:FKBP51:p23 complex, revealing the basis for FKBP51 recognition of this conformation. With these findings we propose a model in which FKBP51 acts specifically during the client maturation step, binding Hsp90 in the ATP-bound state following client loading and NTD dimerization (Figure 6C). From the structure we identify that the TPR H7e helix from FKBP51 binds across a hydrophobic groove in the Hsp90 CTD dimer, which appears distinctly accessible in the closed state compared to the apo Hsp90 CTD (Figure 4). Interactions by the TPR with the Hsp90 CTD enable FKBP51 to extend along the Hsp90 MD with its PPIase-active FK1 domain positioned adjacent Hsp90 client binding loops, potentially enabling FKBP51 catalytic activity to be directed at specific proline residues in the bound client.

Recent cryo-EM structures of the client-bound maturation state reveal Hsp90 catalyzes dramatic client unfolding, likely upon NTD dimerization and formation of the closed conformation (Noddings et al., 2020; Verba et al., 2016). Notably, the N-lobe of Cdk4 is unfolded and threaded though the MD dimer cleft in the Hsp90:Cdc37:Cdk4 structure exposing a strand of residues that would be inaccessible in the folded state (Verba et al., 2016). Thus, the position of the FK1 domain in our structure appears optimal for catalyzing isomerization of specific proline residues that may be localized in unfolded regions or adjacent sites that are optimally presented in the maturation state. Moreover, FKBP51 may accelerate client folding on Hsp90 or stabilize an activated state that is competent for ligand binding or other downstream function. Indeed, modeling of Cdk4 into the Hsp90:FKBP51:p23 structure reveals that P173, a recently-identified FKBP51 target (Ruiz-Estevez et al., 2018), could dock into the FK1 active site following minor FKBP51 conformational changes (Figure S6B).

Hsp90:FKBP51 functions in tau proteostasis, increasing its stability and altering phosphorylation states (Blair et al., 2013; Jinwal et al., 2010). By NMR, Hsp90 is identified to interact with tau transiently in the apo state and function as a scaffold in complex with FKBP51 (Karagoz et al., 2014; Oroz et al., 2018). While Hsp90-FKBP51 contacts we identify in the cryo-EM structure appear similar to CTD and MD interaction regions identified by NMR, we observe only weak binding to the apo state by SEC. Additionally, contrary to the apo state interaction, our data and cryo-EM structure establish that Hsp90 binds FKBP51 with a specific 2:1 stoichiometry that is defined by the H7e binding across the CTD dimer interface. This binding groove can accommodate a single H7e in the closed state, thereby excluding formation of a tetramer complex. Moreover, in the apo state the CTD dimer cleft has reduced accessibility and may be incompatible for H7e binding. Thus, for a reduced affinity apo interaction we postulate that the H7e is likely disengaged, resulting in FKBP51 being more flexibly bound through the EEVD contact alone, in agreement with the dynamic interactions observed by NMR. Alternatively, engagement by the H7e may induce widening of the CTD, enabling its accommodation. However, in existing closed state structures (Huck et al., 2017; Verba et al., 2016) the CTD adopts this widened conformation in the absence of a bound H7e moiety, indicating this may result from NTD dimerization alone.

The residues in H7e identified to contact the Hsp90 CTD groove (notably Y409, M412, and F413) are conserved among TPR-containing PPIase cochaperones (FKBP51, FKBP52, FKBP36, FKBP38 and Cyp40) and part of a region previously designated as the ‘Charge-Y’ motif that was shown to be important for Hsp90 binding (Figure S5F) (Cheung-Flynn et al., 2003). FKBP51/52 are highly similar in their structure and sequence (60% sequence identity) (Wu et al., 2004). Both are found in GR maturation complexes and appear to have opposing roles in dynein-mediated nuclear localization (Grad and Picard, 2007; Wochnik et al., 2005). Only minor differences between FKBP52 and FKBP51 are identified in the crystal structures due to changes in the FK2-TPR domain orientation (Hahle et al., 2019; Wu et al., 2004). This could lead to different positioning of the FK1 domain when bound to Hsp90. However, the FK domains are connected by linkers and based on the cryo-EM density, appear flexible when bound to Hsp90. Thus, we postulate that the organization of FKBP51/52 on Hsp90 is highly similar but with modestly different positioning of the FK1 domains that may be further specified by additional client and cochaperone interactions. Indeed, the proline-rich loop (residues 116-124) in of FKBP51 and FKBP52 is proposed to recognize different conformations of GR in regulating steroid hormone binding (Riggs et al., 2007) and may contribute to conformational differences. Conversely, Cyp40 and FKBP38 are more structurally diverse, containing single FK-like or Cyp (cyclosporin-binding) PPIase domain, respectively, and therefore likely interact differently with Hsp90. Nonetheless, based on the well-defined TPR contacts we identify, Hsp90 interactions with the PPIase cochaperones are expected to be largely conserved and defined by the TPR-CTD interaction.

The Hsp90-p23 contacts appear to be largely conserved between yeast and human Hsp90 systems. However, unlike the crystal structure of yeast Hsp90:p23, the structures presented here (Hsp90:FKBP51:23 and Hsp90:p23) as well as the recent cryo-EM structure of GR:Hsp90:p23 (Noddings et al., 2020), indicate a single p23 molecule appears to bind the human Hsp90 dimer in the maturation state. In our classification and refinements, density for p23 appears on the opposite side of Hsp90 from FKBP51. However, the FKBP51 interaction site does not overlap substantially, indicating p23 binding may only slightly favor the opposite NTD interface and in one class partial density for p23 was resolved on the same side as FKBP51 (Figure S4E). In the GR:Hsp90:p23 structure (Noddings et al., 2020), p23 directly contacts the GR client and enhances ligand binding via its C-terminal tail helix, which is unresolved in our structures. Thus, the positioning of p23 and FKBP51 are likely further specified by interactions with the bound client.

The TPR:EEVD interaction is known to provide the majority of binding energy for Hsp70/Hsp90:cochaperone complexes, enabling cochaperones such as PP5, Hop and CHIP to function in both Hsp70 and Hsp90 pathways (Assimon et al., 2015; Connarn et al., 2014; Scheufler et al., 2000; Smith et al., 2013). The ∼30 residue linker connecting the EEVD to the Hsp90 CTD is not resolved in the Hsp90:FKBP51:p23 structure, indicating these residues do not contribute to FKBP51 binding. Hsp70 similarly contains an unstructured EEVD-connecting linker that may not participate directly in cochaperone interactions (Zhang et al., 2015). These linkers, therefore, may primarily serve to enable TPR-cochaperones to be positioned at different binding sites on Hsp90 or Hsp70 in facilitating their diverse functions.

Our mutagenesis data of H7e indicate this element is critical for Hsp90 binding, and the TPR:EEVD interaction alone is insufficient for stable formation of the complex (Figure 4D). These data, along with the SEC-MALS analysis of open and closed Hsp90 states and position of H7e identified in the structure, reveal that the H7e functions as a key recognition element that enables FKBP51 to bind Hsp90 over Hsp70 and, further, interact specifically with the closed state conformation. Thus, these additional cochaperone:chaperone binding interfaces revealed in structures of intact complexes are likely necessary for enabling cochaperones to engage different steps of the chaperone cycle and client-folding states. Indeed, structures of Hsp90:Hop and, recently, GR:Hsp90:Hsp70:Hop identify that Hop stabilizes the Hsp90 client loading state and participates in client interactions, revealing an active role for Hop in the folding cycle beyond tethering Hsp70 and Hsp90 during client hand-off (Southworth and Agard, 2011; Wang et al., 2020).

## Acknowledgments

We thank K. Lopez, A. Rizo and D. Agard for feedback on the manuscript. We thank the UCSF BACEM Facility for assistance with data collection. This work was supported by an Alzheimer’s Association Research Fellowship (to K.L.), and NIH grants AG002132 and AG068125 (to D.R.S.).

## Author Contributions

K.L. performed biochemical and cryo-EM experiments, developed figures and wrote and edited the manuscript. A.C.T. generated mutant constructs and expressed and purified proteins. E.T. operated the Krios microscope and helped with data collection and structure determination. S.N.G. performed initial biochemical and cryo-EM analysis and edited the manuscript. D.R.S. designed and supervised the project and wrote and edited the manuscript.

## Declaration of Interests

The authors declare no competing interests.

## Supplemental Information

**Figure S1.**
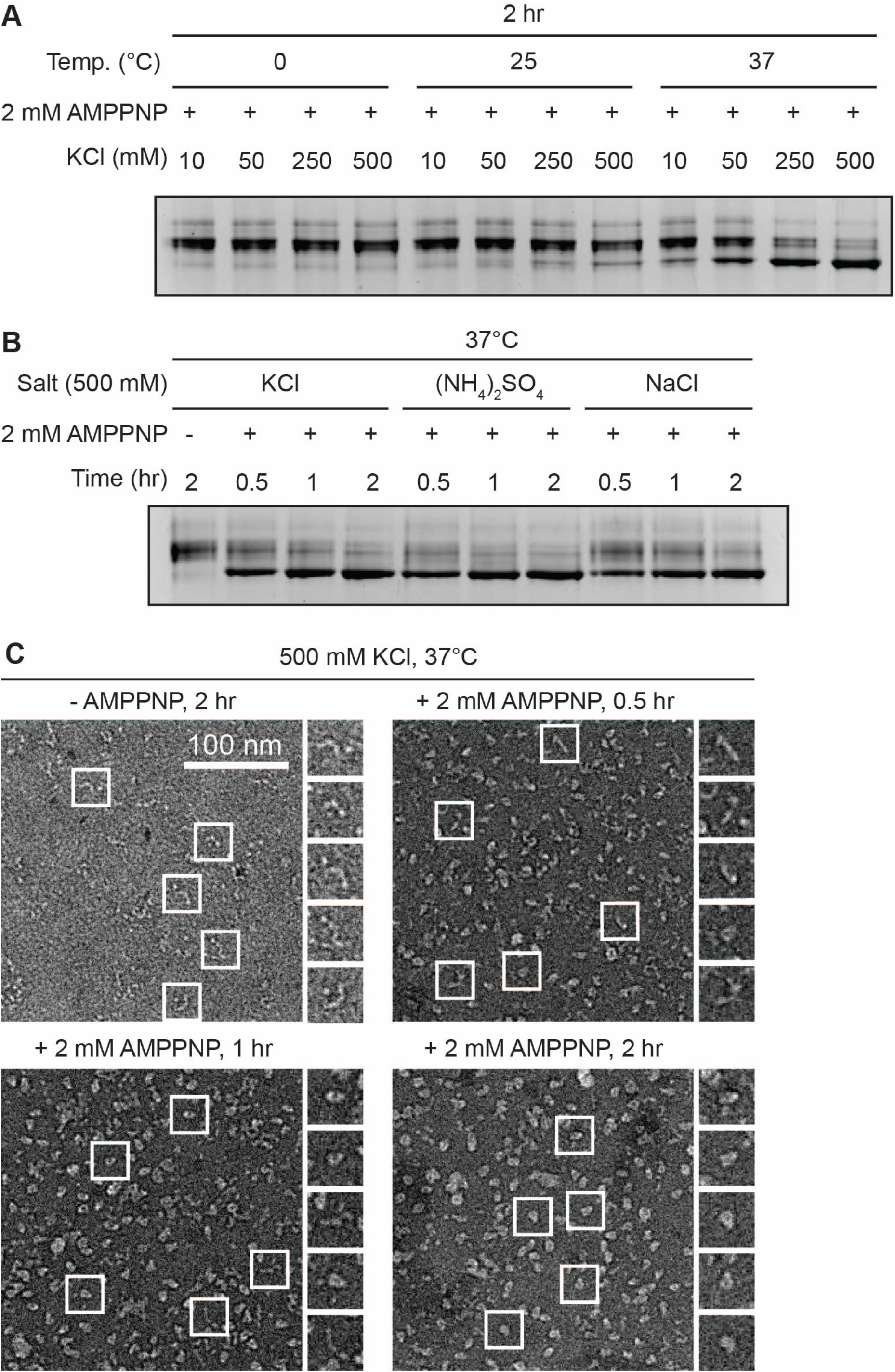
Formation of the closed, ATP state of Hsp90. (A) Native-gel analysis of the Hsp90 dimer following incubation at indicated temperature and KCl concentrations, and (B) with different salts. Higher mobility band is indicative of the closed, ATP conformation of the dimer and increases with increasing salt concentration and temperature. (C) Representative negative-stain EM images of Hsp90 following incubation at indicated conditions, showing formation of the closed conformation following 2-hour incubation at 37°C with 500 mM KCl and 2 mM AMPPNP.

**Figure S2.**
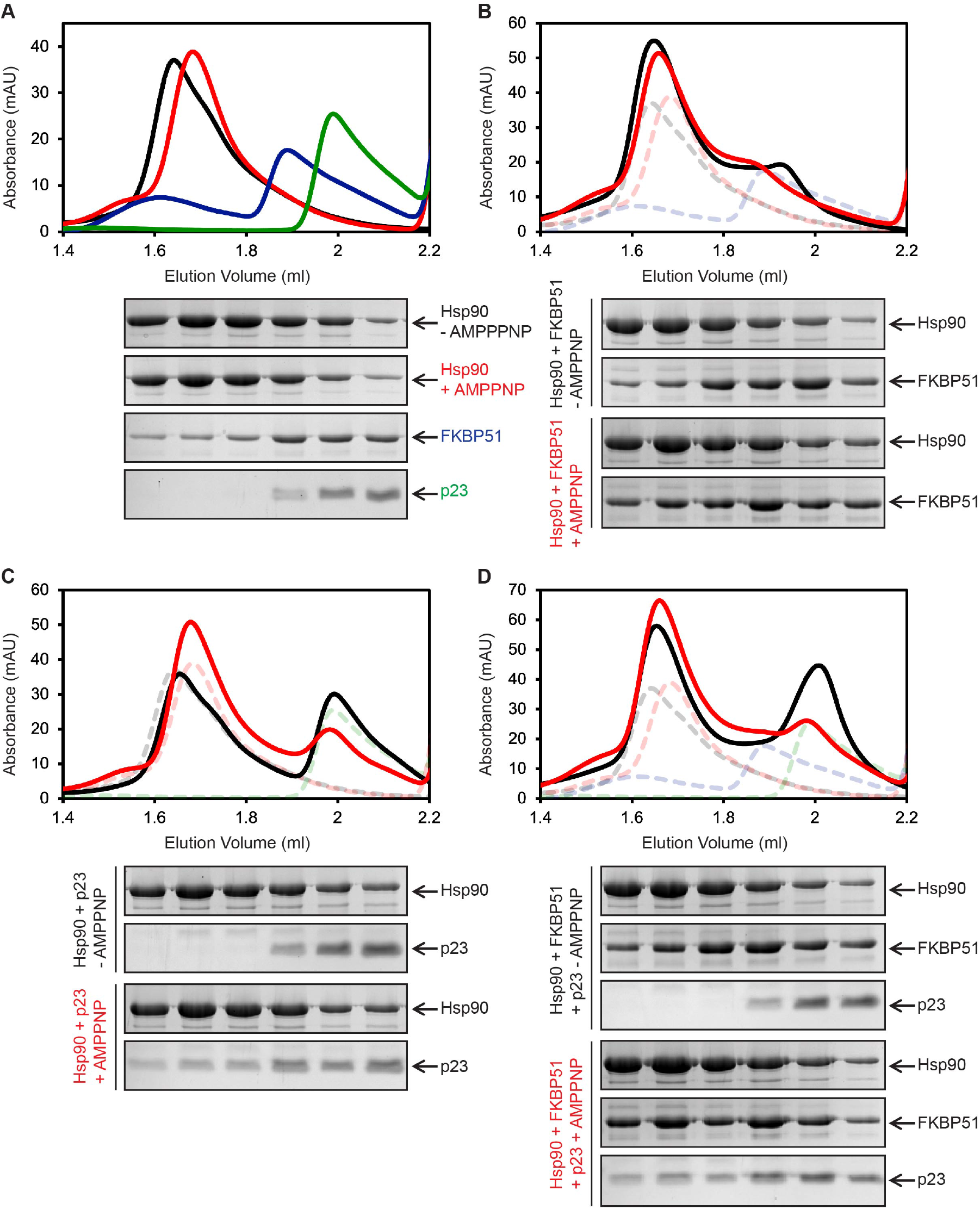
SEC and gel analysis of Hsp90:FKBP51:p23 complex formation for the open and closed Hsp90 states. Hsp90 was incubated with 500 mM of KCl in the absence (black) or presence (red) of 2 mM AMPPNP at 37°C for 2 hours (to form the open and closed states, respectively) (A) alone, or with (B) FKBP51, (C) p23, or (D) both FKBP51 and p23. Dashed line traces represent runs for the individual proteins: Hsp90-open (black), Hsp90-closed (red), FKBP51 (blue), and p23 (green). SDS-PAGE analysis of the SEC fractions is shown beneath the chromatogram with proteins indicated and each lane aligned to the elution volume. Note: a minor oligomeric form of FKBP51 elutes at 1.6 ml (blue trace in A) and contributes to the presence of FKBP51 in early fractions with Hsp90 in the open state (B and D). However, by SEC-MALS analysis (Figure 2) the mw_avg_ indicates Hsp90 elutes primarily as a dimer and is unbound to FKBP51 under these conditions.

**Figure S3.**
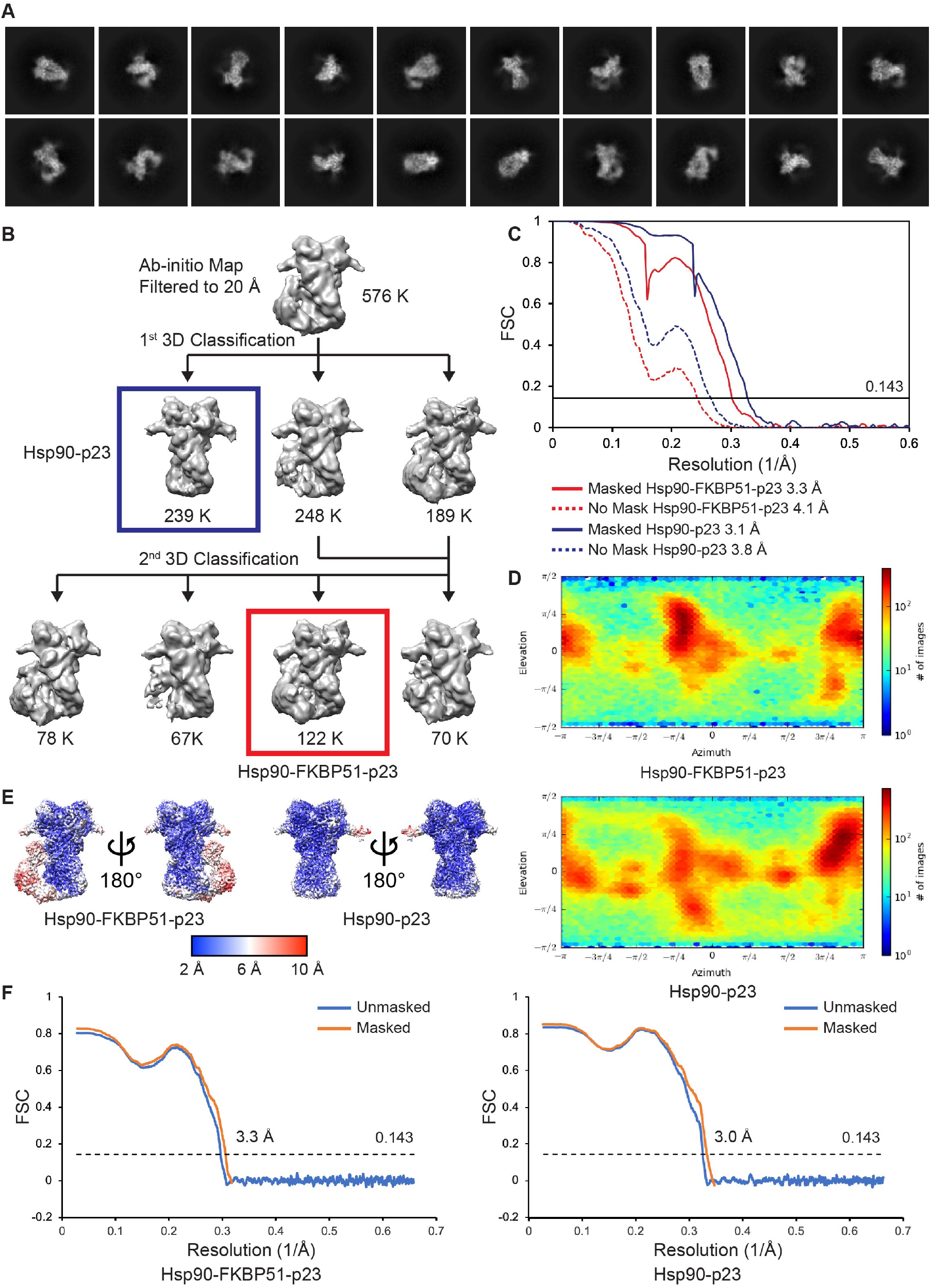
Cryo-EM data processing and structure determination procedure. (A) Representative reference-free 2D class averages of Hsp90:FKBP51:p23 particles (Box size equals 300 Å). (B) Cryo-EM 3D classification scheme to achieve classes used for refinement processing of Hsp90:FKBP51:p23 (red) and Hsp90:p23 (blue) structures. (C) Gold standard Fourier shell correlation (FSC) curves of the final masked and unmasked refinements of Hsp90:FKBP51:p23 and Hsp90:p23 complexes. (D) 3D angular distribution plots of the particles used in the final reconstruction (E) Cryo-EM maps of the final Hsp90:FKBP51:p23 and Hsp90:p23 reconstructions colored by local resolution, determined using cryoSPARC2 (Punjani et al., 2017). (F) Map vs. Model FSCs of Hsp90:FKBP51:p23 and Hsp90:p23 complexes.

**Figure S4.**
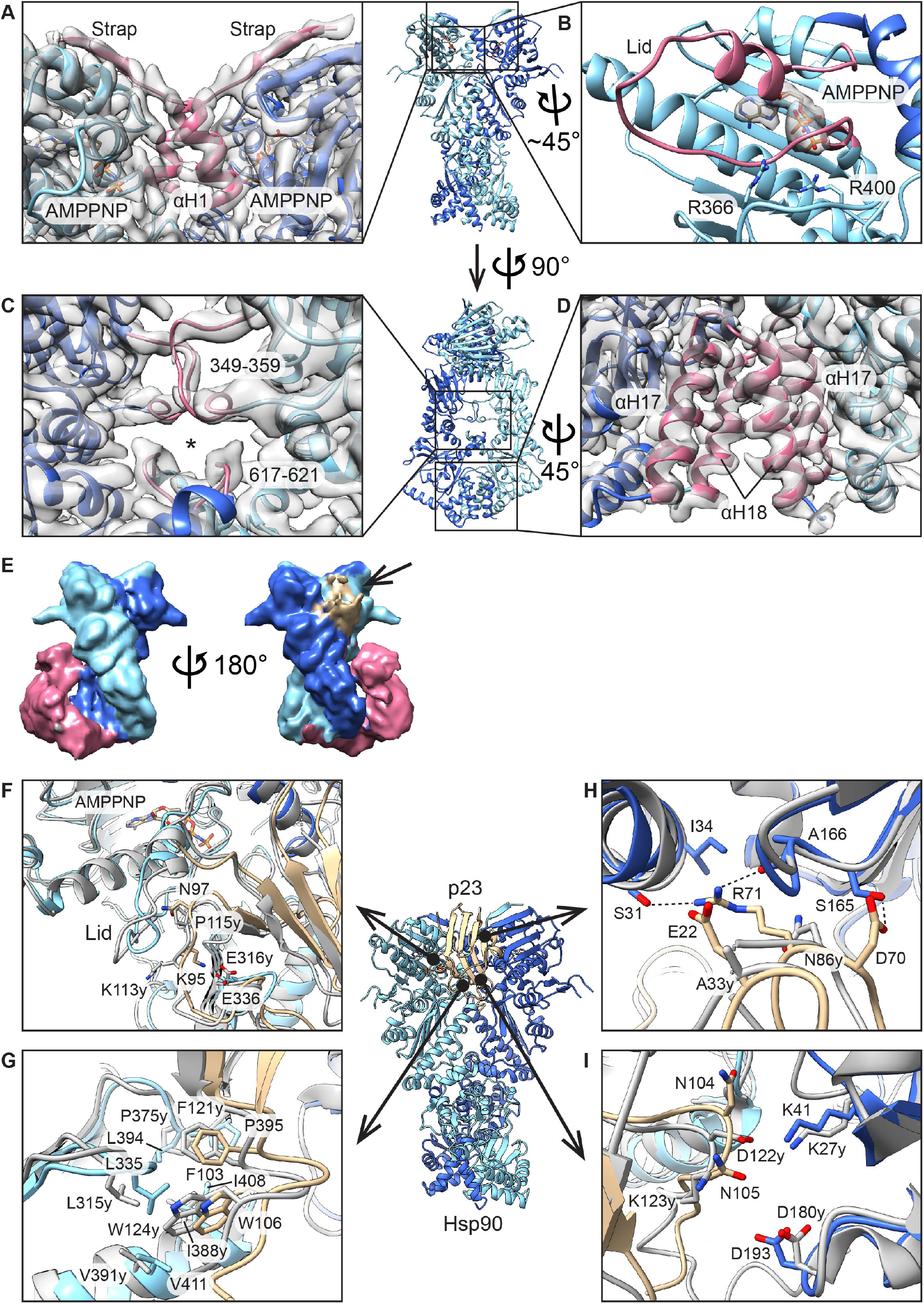
The cryo-EM maps and the models of Hsp90:FKBP51:p23 and Hsp90:p23. (A-D) Structural features of Hsp90 (A) The N-terminal straps (residues 17-15, red) form beta sheet structures across the top of the NTDs and αH1 from each protomer interacts each other (red). (B) The segmented map showing a density for AMPPNP in the NTD. The α-helical lid segment (residues 109-139, red) encompasses AMPPNP at the nucleotide pocket. R400 in the middle domain coordinates γ-phosphate of AMPPNP and R366 closes the lid, acting as a latch. (C) The loops in the middle domain (residues 349-359) and the C-terminal domain (residues 617-621) involved in substrate interaction are shown in red. These loops form a mostly hydrophobic ‘tunnel’ (asterisk) that captures threaded substrates (Verba et al., 2016). (D) αH17 and αH18 of each protomer form C-terminal dimerization interface (red). (E) The cryo-EM map of class 1 after second round of 3D classification shown in Figure S3B. Densities are colored based on fit of model determined from the Hsp90:FKBP51:p23 structure but with partial density for p23 present on the same side as FKBP51. Densities correspond to each monomer of Hsp90 (light blue and blue), FKBP51 (pink), and p23 (tan). Partial density for p23 is indicated (arrow). (F-I) Conserved Hsp90-p23 interactions in the closed, ATP state. The Hsp90:p23 model determined here and colored as in Figure 3 is aligned to the yeast Hsp90:p23 crystal structure (PDB: 2CG9, grey) (Ali et al., 2006). (F) N97 (P115 in yeast) of p23 interacts with the lid of Hsp90. K95 of p23 forms a salt bridge with E336 of Hsp90 in the middle domain. (G) Conserved F103 and W106 of p23 interact with the hydrophobic pocket of Hsp90, consisting of L335, L394, P395, I408, and V411 at the middle domain. (H) D70 and R71 of p23 form a hydrogen bond network with the N-terminal domain (S31, A166, and S165) of Hsp90, indicated with dashed line. This network is not conserved in yeast. (I) Salt bridges identified in the yeast structure (D122-K27 and K123-D180) are not observed in human.

**Figure S5.**
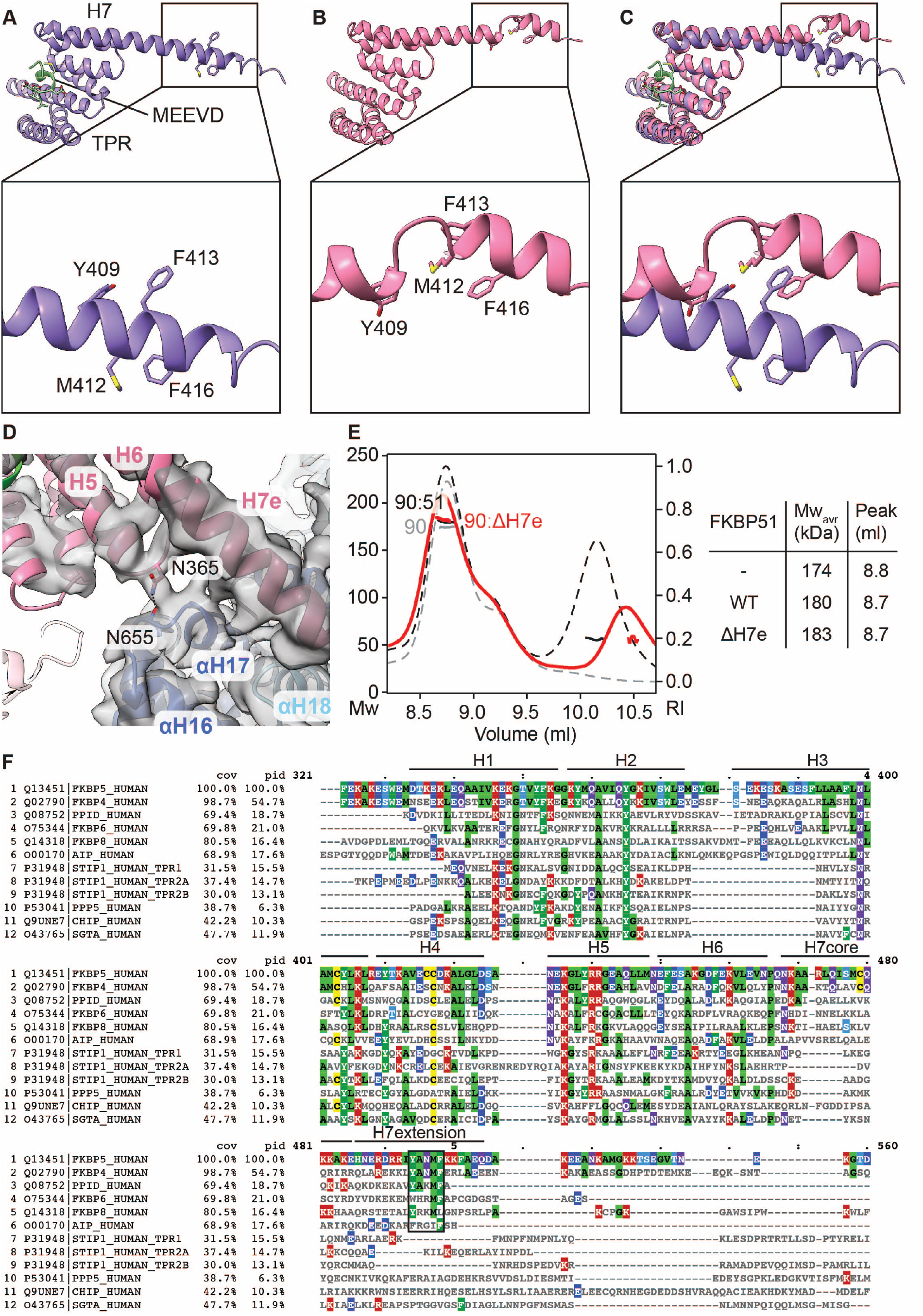
Characterization of the FKBP1 TPR domain helix 7 extension (H7e). (A-C) Comparison of the H7e conformation from the crystal structure (PDB: 5NJX) (Kumar et al., 2017). (A) and the Hsp90:FKBP51:p23 structure determined here (B) showing partial unfolding of the helix at positions 409-416, which contacts the CTD dimer interface in Hsp90. (D) Map+model view of the FKBP51 TPR-Hsp90 interaction showing contact between FKBP51 N365 and Hsp90 N665. (E) SEC-MALS analysis of FKBP51 ΔH7e (Δ401-457) binding to Hsp90 in the open state, without 2 mM AMPPNP (red), showing no significant interaction, as observed with wildtype (black) and in comparison to Hsp90 alone (grey). (F) Alignment of the TPR domain sequences for immunophilins: FKBP5, FKBP4, PPID, FKBP6, FKBP8, and AIP and cochaperones: STIP1, PPP5, CHIP, and SGTA. The TPR domain helixes are numbered and conserved hydrophobic residues in the H7extension involved in the Hsp90 CTD interaction are indicated (black rectangle).

**Figure S6.**
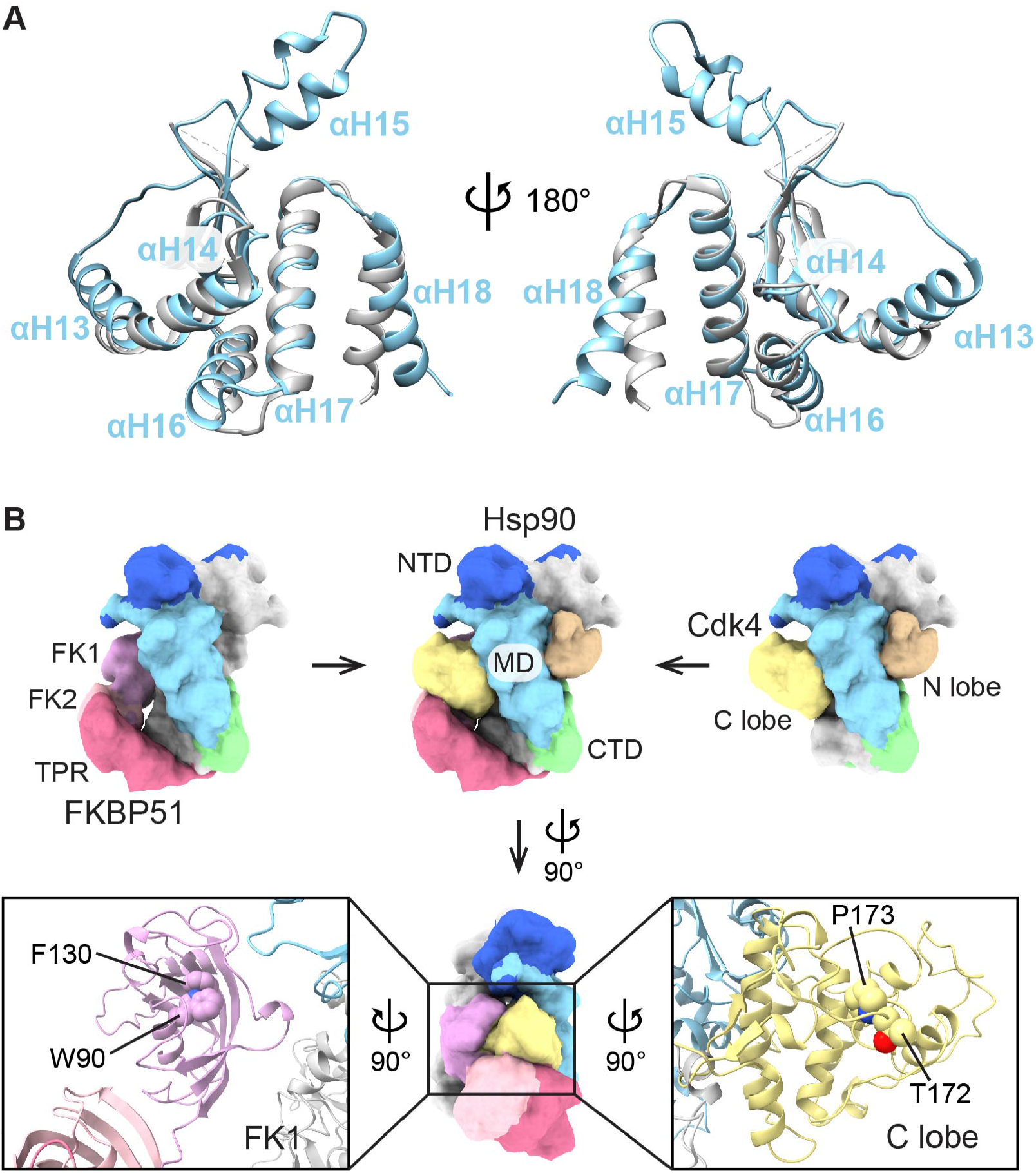
Comparison of the C-terminal domain of Hsp90 in different states and a model of Cdk4 regulation by FKBP51 coordinated by Hsp90. (A) The open, apo (grey) state of Hsp90 was modeled based on E. coli Hsp90 open structure (PDB: 2IOQ) (Shiau et al., 2006) using SWISS-MODEL (Waterhouse et al., 2018). The model and the closed, ATP (light blue) state structure were aligned at αH17 using Chimera. (B) Each domain of FKBP51, Hsp90, and Cdk4 are indicated with domain name and unique color (top). Cdk4 position relative to Hsp90 is adopted from previous cryo-EM structure of Hsp90:Cdc37:Cdk4 (PDB: 5FWK) (Verba et al., 2016). Cryo-EM structure of Hsp90:FKBP51 complex (left) and cryo-EM structure of Hsp90:Cdk4 complex are superimposed based on Hsp90 (middle). P173 in the C lobe (bottom, right) of Cdk4 might be specifically isomerized by FK1 domain (bottom, left). Amino acids (W90 and F130) comprising the active site are shown in sphere representation. Phosphorylation of T172 of Cdk4 regulates isomerization of P173 (Ruiz-Estevez et al., 2018).

## Notes

### Competing Interest Statement

The authors have declared no competing interest.

